# Classifying COVID-19 variants based on genetic sequences using deep learning models

**DOI:** 10.1101/2021.06.29.450335

**Authors:** Sayantani Basu, Roy H. Campbell

## Abstract

The COrona VIrus Disease (COVID-19) pandemic led to the occurrence of several variants with time. This has led to an increased importance of understanding sequence data related to COVID-19. In this chapter, we propose an alignment-free k-mer based LSTM (Long Short-Term Memory) deep learning model that can classify 20 different variants of COVID-19. We handle the class imbalance problem by sampling a fixed number of sequences for each class label. We handle the vanishing gradient problem in LSTMs arising from long sequences by dividing the sequence into fixed lengths and obtaining results on individual runs. Our results show that one-vs-all classifiers have test accuracies as high as 92.5% with tuned hyperparameters compared to the multi-class classifier model. Our experiments show higher overall accuracies for B.1.1.214, B.1.177.21, B.1.1.7, B.1.526, and P.1 on the one-vs-all classifiers, suggesting the presence of distinct mutations in these variants. Our results show that embedding vector size and batch sizes have insignificant improvement in accuracies, but changing from 2-mers to 3-mers mostly improves accuracies. We also studied individual runs which show that most accuracies improved after the 20th run, indicating that these sequence positions may have more contributions to distinguishing among different COVID-19 variants.

## 1.1 COVID-19 and Genetic Data

The COrona VIrus Disease (COVID-19) resulted in more than 170 million cases and more than 3 million deaths worldwide [1]. The virus causing this disease is known as the SARS-CoV2 virus with the first recorded cases in 2019 [2]. In addition to the burden on healthcare resources, other impacts were also observed in other industries including education, economy, and travel [3, 4]. Countries all around the world tried to curb the infections by employing a series of mitigation measures in the form of lockdowns, face coverings, social distancing, frequent sanitizing, limits on gatherings, etc [5]. There have been several research efforts worldwide to propose and design drugs for treating COVID-19 [6, 7]. In severely affected COVID-19 patients, ventilators and antipyretics have shown varying degrees of efficacy [8, 9].

The common “end” to pandemics is achieved through herd immunity, which can be achieved by ensuring that a significant (preferably the entire) population has antibodies in their system [10]. Vaccination is one such way to achieve this. In December 2020, several vaccine candidates were approved after clinical trials in an effort to protect the population from COVID-19 [11]. However, due to the virulent nature of COVID-19, mass vaccination with careful mitigation measures still need to be employed [12] until significant portions of the population including the vulnerable groups have been fully vaccinated. Studies are still ongoing about the duration of protection provided by these vaccines and whether booster shots will be required for further protection [13]. Another problem that has surfaced with time is the rise of COVID-19 “variants”, which are mutations of the original virus that may or may not be more infectious [14]. Most trials show that most COVID-19 vaccines are effective against such variants, although further studies are needed in order to study the newer variants with time [15, 16, 17]. The most important aspect in drug design and vaccine design is understanding the underlying make-up of the virus – this includes studying mutations occurring in all variants of the virus. As a result, sequence data is essential in an effort to fully understand the nature of the COVID-19 virus.

Machine learning, specifically deep learning has been helpful in understanding and modeling the COVID-19 pandemic with the enormous amount of data available — several studies have been carried out on image data [18, 19, 20], epidemiological data [21, 22, 23, 24], and sequence data [25, 26, 27]. At this point of the pandemic, the rise of several COVID-19 variants has led to an increased need of studying genetic sequences. More importantly, a system to automatically distinguish and classify the COVID-19 variants using deep learning will help contribute to ongoing research efforts along with vaccine and drug development research.

In this chapter, we propose an LSTM (Long Short-Term Memory) [28] based deep learning model to classify different COVID-19 variants based on genetic sequence data. The rest of this chapter is organized as follows: Section 1.2 discusses the various COVID-19 variants being studied, Section 1.3 presents our data and methods, Section 1.4 discusses our results, Section 1.5 concludes this chapter, and Section 1.6 mentions our code repository and availability of our results.

## 1.2 COVID-19 Variants

In this section, we discuss the various COVID-19 variants which have been used in our experiments. We discuss all variants studied with regard to the PANGO lineage convention in the following subsections.

### 1.2.1 B.1.1.214

This variant was observed in Japan during the third COVID-19 wave [29] which has been identified to be of 501N+484E type [30].

### 1.2.2 B.1.1.519

This variant was identified as a variant of interest (VOI) in Mexico [31]. Rhoads et al. [32] have observed the g.29197C>T mutation in all cases of this variant.

### 1.2.3 B.1.160

This variant predominantly appeared during the second COVID-19 wave in Eastern Germany, prior to which it was observed in several other European countries [33].

### 1.2.4 B.1.177.21

This variant was found to have spread in several European countries [34].

### 1.2.5 B.1.177

This is the parent lineage of variant B.1.177.21 [34]. The parent variant is more widespread and has several types of strains [35].

### 1.2.6 B.1.1.7

Also known as the Alpha variant, this variant is widespread and infectious [36] and has affected several countries including the United States [37].

### 1.2.7 B.1.1

This is another variant of COVID-19 that has been detected in several countries [38].

### 1.2.8 B.1.221

This variant of COVID-19 was one of the variants detected predominantly in Eastern Germany leading to the second wave as studied by Yi et al. [33].

### 1.2.9 B.1.243

This variant was a variant of interest (VOI) that emerged in Arizona, United States containing the E484K mutation and spread to other parts of the United States [39].

### 1.2.10 B.1.258

This variant was observed to spread in several countries including the Romania [40], East Germany [33], Lebanon [41], Czech Republic and Slovakia [42].

### 1.2.11 B.1.2

This variant of COVID-19 was observed to have emerged in Brazil [43].

### 1.2.12 B.1.351

Also known as the Beta variant, this widespread and infectious variant [36] is known to have the E484K, K417N, N501Y substitutions [37].

### 1.2.13 B.1.427

Also known as the Epsilon variant, this COVID-19 variant was identified in parts of the United States and was found to be not as infectious as the Alpha, Beta, and Gamma variants [44, 45].

### 1.2.14 B.1.429

Also known as the Epsilon variant, this variant also belongs to the same lineage as that of B.1.427 [45].

### 1.2.15 B.1.526

Also known as the Iota variant, this variant was identified in New York, United States [46]. It contains the E484K mutation [47].

### 1.2.16 B.1.596

This variant of COVID-19 was observed to have spread in several countries world-wide [48].

### 1.2.17 B.1.617.2

Also known as the Delta variant, this variant of COVID-19 was observed to have contributed to the surges of the pandemic in India [49] and has also been observed in England [50] and the United States [37].

### 1.2.18 B.1

This is the variant of COVID-19 that constitutes the parent lineage of several COVID-19 variants circulating across the various countries around the world [51].

### 1.2.19 D.2

This is the variant of COVID-19 that has been predominantly observed in Australia [52].

### 1.2.20 P.1

Also known as the Gamma variant, this variant was first observed in Brazil [53] and has affected several countries worldwide including the United States [37].

## 1.3 Data and Methods

Data for this study was collected from GISAID (https://www.gisaid.org). We collected labeled data from 20 lineages of COVID-19 containing genetic sequences in FASTA format from the beginning of the COVID-19 pandemic till June 5, 2021. For the purpose of this study, unclassified sequences that were part of ‘None’ were not considered.

All experiments in this study were run on `gpux1` on the HAL (Hardware-Accelerated Learning) cluster [54].

The presence of multiple variants in the dataset included some variants that were more widespread and virulent than others and some variants that had more genetic samples collected in general. However, from a machine learning perspective, training on samples with imbalanced labels would lead to a class imbalance problem [55]. In order to handle this, we decided to use sampling prior to training our model. For experiments in this study, we limit the number of labels per class to 1000. The sampled sequences were chosen randomly from each of the lineages using random.choice from numpy [56].

We have designed our experiments as follows:

1. One-vs-all classification: In this part of our study, we trained and tested our LSTM model on 20 different binary classifiers. Each classifier was trained to identify a specific variant from the rest. As previously discussed, for each classifier, we handle the class imbalance problem in this scenario by considering 1000 sequences from a specific class and 1000 sequences sampled from the rest of the classes.
2. Multi-class classification: In this part of our study, we trained and tested our LSTM model to classify among 20 different variants at once using a single classifier. As previously discussed, in this case too, we handle the class imbalance problem by considering 1000 sequences that are sampled for each class label.

We propose an LSTM (Long Short-Term Memory) [28] based model that would classify COVID-19 sequences into their respective variants. LSTMs are a type of recurrent neural network model used in deep learning that are capable of capturing dependencies over long periods of time and can be used for modeling time series data. This is useful because variants have temporal consistency making them more obvious to an LSTM model. Each gene sequence is divided into consecutive overlapping k-mers. The k-mers when counted uniquely constitute the vocabulary of the language being considered. For all DNA sequences, we consider the entire alphabet, that is, *{A,T,G,C}* along with the ambiguous nucleotides – *{S,W,R,Y,K,M,B,D,H,V,N}*. A distribution of k-mers revealed the more common nucleotides are *{A,T,G,C,N}*. We have included all unambiguous as well as ambiguous nucleotides since the number of ambiguous nucleotides was small enough compared to the unambiguous nucleotides.

In essence, every DNA sequence is transformed into a set of ‘words’ and we study the ordering of these words using an LSTM based model in order to classify the sequences into their respective variants or classes. We used integer encoding and padding prior to feeding in the sequences into our LSTM framework, all coded using keras [57] with a Tensorflow [58] backend in Python3. However, a problem with directly feeding in sequences in this manner is that in designing such an LSTM, the sequence length, and consequently, the integer encoded words are essentially the timesteps of the model. The minimum sequence length observed in our studies was around 26000. We modeled this as several timesteps in an LSTM model because of size and to prevent the vanishing gradient problem [59]. Moreover, experiments where we directly attempted to train the LSTM model with the full sequences at once were slow to run due to excessive computational overhead and the programs eventually did not execute until completion. In an effort to overcome this, we divided the problem into 51 steps, where we feed in segments of 500 nucleotides (characters of the genetic sequences) at a time. This helped the programs run successfully until completion. Each of the predictions from the test set were then averaged over the 51 steps and thresholded with 0.5 in order to obtain the overall accuracy of the model. This framework is suitable for the following reasons: (1) it helps reduce computational overload of feeding in multiple timesteps at once, (2) it handles the vanishing gradient problem of LSTMs that can arise due to a large number of timesteps, and (3) it enables us to find distinguishing portions of the genetic sequences that contribute to better accuracies — such portions of data are “useful” for identifying and classifying the specific COVID-19 variant. The only problem with this approach would be missing nucleotides of lengthy sequences located towards the end due to truncating the sequences after fixed indices in order to allow the model to train effectively. It is also to be noted at this point that we have used an alignment-free k-mer based approach in all our experiments in this study.

### 1.3.1 Model Architecture

The value of *k* for generating k-mers is a hyperparameter itself. Small k-mer values help in faster extraction of k-mers from long sequences, but do not provide adequate coverage compared to larger k values.

For our proposed framework, we use an embedding layer to convert the integer encoded k-mer words into vectors of a fixed size. The embedding vector size constitutes a hyperparameter, along with the batch size that we varied in our experiments.

We use 10 LSTM units and 1 dense layer with sigmoid activation in order to obtain the probabilities in the case of all our binary classification one-vs-all models. We use Adam [60] as the optimizer with binary cross entropy and accuracy as the metric in order to train the model for 50 epochs. We use Sequential() from keras to stack the layers in a linear fashion.

We used 10 LSTM units and 20 dense units with softmax activation for our multi-class classification models. We use Adam [60] as the optimizer with sparse categorical cross entropy and accuracy as the metric in order to train the model for 50 epochs. Similar to the one-vs-all classifiers, we use Sequential() from keras to stack the layers in a linear fashion.

## 1.4 Results and Discussion

The entire set of sampled sequences was shuffled retaining original variant labels and then divided into an 80% and 20% ratio for training and testing data respectively. In the results, we primarily focus on discussing accuracies on test data since these comprise results on “unseen” data and are representative of what the LSTM model has learnt from the training process.

Table 1.1 shows the accuracies of varying hyperparameters for k=2. The cells in the table show accuracy as a percentage calculated as a combination of the predictions across all 51 runs. The predicted probabilities from the Dense layer are thresholded using 0.5 to represent whether a sample belongs to a class or not. The predictions are then compared with the ground truth class labels and the accuracies for every classifier are obtained using sklearn [61]. Each row in the table represents a classifier where the task is classifying the specific variant from other COVID-19 variants. Based on our experiments, changing the embedding size and batch size did not have significant effect on the accuracies. It is important to note that certain classifiers like those of B.1.1.214, B.1.177.21, B.1.1.7, B.1.526, and P.1 have better accuracies compared to others – these higher accuracy scores may be because we are essentially trying to classify one specific variant and it would need to have features that are distinct enough for the classifier to tell it apart from the other classes. This may be a broader indicator of variants that have “novel mutations” of COVID-19 with genetic makeup different enough from the rest of the sequences from other variants.

**Table 1.1.**
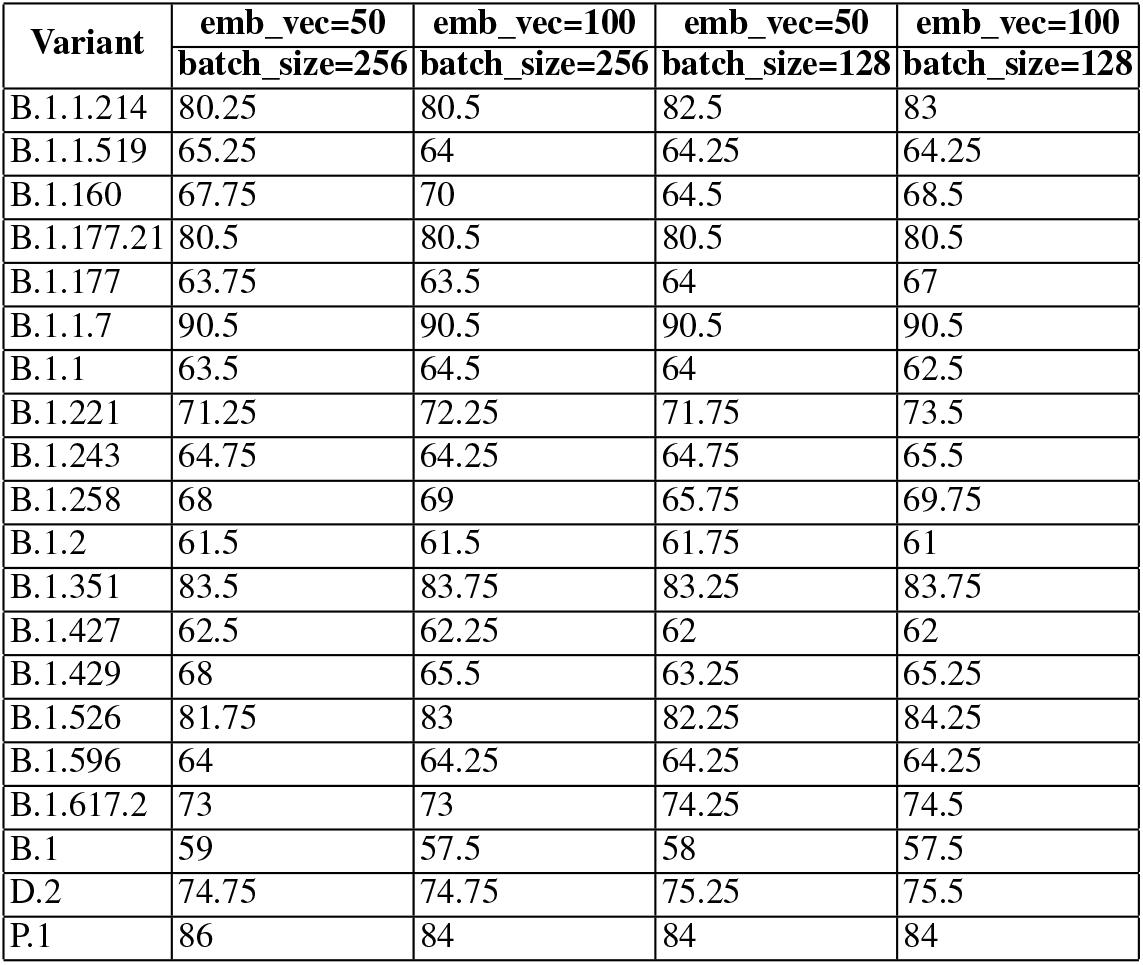
Accuracies for classifying COVID-19 lineages with k=2

Within the accuracies from individual runs, positions after the 20th run till the 51st run showed higher accuracies in general, hence indicating that these positions may be of more interest in contributing to the “distinction” of genetic makeup compared to the beginning portions of the sequences. The exact positions vary based on the specific classifier and hyperparameters. The entire sets of accuracies obtained from our individual runs as well as the final results are included in our code repository.

Table 1.2 shows the accuracies of varying hyperparameters for k=3. While there is not much significant change in accuracies compared to the experiments for k=2, it is interesting to note that for certain cases, it is beneficial to use k=3. In this case too, changes in embedding vector size and batch size did not lead to any significant changes in the accuracies. The most significant increase in accuracy was observed in the case of B.1.160. This may be because 3-mers may be more representative of distinguishing characteristic features of this variant compared to 2-mers. Based on the results, accuracies generally improve after the 20th run which indicates that these positions may be of interest in distinguishing characteristic features for a certain class similar to the 2-mers. It is to be noted that while increasing the value of k in k-mers may add more coverage, the process of extracting sequences takes longer time.

**Table 1.2.** Accuracies for classifying COVID-19 lineages with k=3

| Variant | emb_vec=50 | emb_vec=100 | emb_vec=50 | emb_vec=100 |
| --- | --- | --- | --- | --- |
|  | batch_size=256 | batch_size=256 | batch_size=128 | batch_size=128 |
| B.1.1.214 | 82 | 81.75 | 82 | 82.5 |
| B.1.1.519 | 63.75 | 63.75 | 64 | 65.5 |
| B.1.160 | 77.75 | 78.25 | 78.25 | 71.5 |
| B.1.177.21 | 80.5 | 80.5 | 80.5 | 80.5 |
| B.1.177 | 67.25 | 66 | 64.5 | 66.75 |
| B.1.1.7 | 91 | 91 | 91 | 92.5 |
| B.1.1 | 63.5 | 63.75 | 63.75 | 63.25 |
| B.1.221 | 73.25 | 73 | 73 | 72.75 |
| B.1.243 | 65 | 65 | 65 | 64.25 |
| B.1.258 | 70.25 | 68.5 | 69.25 | 73.25 |
| B.1.2 | 61.75 | 61.75 | 61.5 | 62.75 |
| B.1.351 | 84 | 84.25 | 84 | 85.5 |
| B.1.427 | 62.25 | 62.25 | 62.25 | 64 |
| B.1.429 | 65 | 65 | 65.75 | 65.75 |
| B.1.526 | 84.75 | 84.5 | 85 | 84 |
| B.1.596 | 65.75 | 65.75 | 66 | 66.5 |
| B.1.617.2 | 74.75 | 74.75 | 74.75 | 75.75 |
| B.1 | 58 | 57 | 57.5 | 56.5 |
| D.2 | 75.75 | 75.75 | 76 | 76.25 |
| P.1 | 84.5 | 84.75 | 84.75 | 85.75 |

For the multi-class classifier, we varied hyperparameters and obtained accuracies for individual runs for embedding vector sizes of 50, 100 and batch sizes of 256, 128 for both k=2 and k=3. The results show accuracies of less than 50% for most of the individual runs in all cases, which are significantly lesser than the accuracies of individual runs obtained for the one-vs-all classifiers. This may be because it is harder to correctly distinguish among 20 classes and classify the same compared to a set of binary classifiers identifying whether a specific variant belongs to a class or not. In terms of individual accuracies of runs, it is more beneficial to build 20 binary classifiers compared to a single classifier classifying 20 classes. While comparing individual accuracies, though the accuracies themselves are low, the accuracies for multi-class classifiers also show an increase in accuracy after the 20th run in general indicating that these may be positions contributing to distinguishing features, which was also observed in case of the one-vs-all classifiers. Results obtained from individual runs of the multi-class classifier are also available in our code repository.

## 1.5 Conclusion

In this chapter, we proposed an LSTM based deep learning model that can classify COVID-19 variants based on genetic sequences. Our proposed method is alignment free and uses k-mers. In our method, we handle the class imbalance problem by sampling a fixed set of sequences per class. We also handle the vanishing gradient problem of LSTMs arising from long timesteps due to long lengths of genetic sequences by dividing the sequences into fixed segments of smaller timesteps. We discuss results for accuracies on both one-vs-all classifiers as well as multi-class classifier models. Our results show that the one-vs-all classifier performs better than the multi-class classifier models for distinguishing variants on individual runs, with test accuracies as high as 92.5%. Higher overall accuracies in variants like B.1.1.214, B.1.177.21, B.1.1.7, B.1.526, and P.1 may suggest that these variants have specific mutations that distinguish them from the other variants in case of the one-vs-all classifiers. In terms of hyperparameters, varying the embedding vector sizes and batch sizes did not change the accuracies in a significant manner based on our experiments. However, k=3 yielded slightly better accuracies compared to k=2 in most cases. We also observed that the accuracies on individual runs improve after the 20th run in general, which may suggest that these positions are more important in determining distinctive features contributing to a specific class. We hope that our model can be extended in the future for understanding COVID-19 variants based on sequence data and believe that it can help contribute to vaccine and drug development research.

## 1.6 Code

Code and results from our experiments are available here: https://github.com/sayantanibasu/covid19-gene-variants.

## Acknowledgements

This project has been funded by the Jump ARCHES endowment through the Health Care Engineering Systems Center.

This work uses resources from GISAID (https://www.gisaid.org). We would like to acknowledge all laboratories that have contributed their COVID-19 sequence data to GISAID.

This work utilizes resources supported by the National Science Foundation’s Major Research Instrumentation program, grant #1725729, as well as the University of Illinois at Urbana-Champaign.

